# Structural basis for neutralization of SARS-CoV-2 and SARS-CoV by a potent therapeutic antibody

**DOI:** 10.1101/2020.06.02.129098

**Authors:** Zhe Lv, Yong-Qiang Deng, Qing Ye, Lei Cao, Chun-Yun Sun, Changfa Fan, Weijin Huang, Shihui Sun, Yao Sun, Ling Zhu, Qi Chen, Nan Wang, Jianhui Nie, Zhen Cui, Dandan Zhu, Neil Shaw, Xiao-Feng Li, Qianqian Li, Liangzhi Xie, Youchun Wang, Zihe Rao, Cheng-Feng Qin, Xiangxi Wang

## Abstract

The COVID-19 pandemic caused by the SARS-CoV-2 virus has resulted in an unprecedented public health crisis. There are no approved vaccines or therapeutics for treating COVID-19. Here we reported a humanized monoclonal antibody, H014, efficiently neutralizes SARS-CoV-2 and SARS-CoV pseudoviruses as well as authentic SARS-CoV-2 at *nM* level by engaging the S receptor binding domain (RBD). Importantly, H014 administration reduced SARS-CoV-2 titers in the infected lungs and prevented pulmonary pathology in hACE2 mouse model. Cryo-EM characterization of the SARS-CoV-2 S trimer in complex with the H014 Fab fragment unveiled a novel conformational epitope, which is only accessible when the RBD is in open conformation. Biochemical, cellular, virological and structural studies demonstrated that H014 prevents attachment of SARS-CoV-2 to its host cell receptors. Epitope analysis of available neutralizing antibodies against SARS-CoV and SARS-CoV-2 uncover broad cross-protective epitopes. Our results highlight a key role for antibody-based therapeutic interventions in the treatment of COVID-19.

**One sentence summary:** A potent neutralizing antibody conferred protection against SARS-CoV-2 in an hACE2 humanized mouse model by sterically blocking the interaction of the virus with its receptor.

## Main Text

The coronavirus disease 2019 (COVID-19) pandemic had afflicted >7 million people in more than 200 countries/regions, resulting in 400,000 deaths as of 8 June, according to the World Health Organization. The etiological agent of this pandemic is the newly emerging coronavirus, severe acute respiratory syndrome coronavirus 2 (SARS-CoV-2), which, together with the closely related SARS-CoV belongs to the lineage B of the genus *Betacoronavirus* in the *Coronaviridae* family (*1*). Sharing an amino acid sequence identity of ∼80% in the envelope-located spike (S) glycoprotein, both SARS-CoV-2 and SARS-CoV utilize human angiotensin converting enzyme 2 (hACE2) to enter into host cells. Cellular entry is achieved by the homotrimeric S mediated virus-receptor engagement through the receptor binding domain (RBD) followed by virus-host membrane fusion (*1, 2*). Abrogation of this crucial role played by the S-protein in the establishment of an infection is the main goal of neutralizing antibodies and the focus of therapeutic interventions as well as vaccine design (*3-6*). Several previously characterized SARS-CoV neutralizing antibodies (NAbs) were demonstrated to exhibit very limited neutralization activities against SARS-CoV-2 (*7-9*). Among these, CR3022, a weakly neutralizing antibody against SARS-CoV, is tight binding, but non-neutralizing for SARS-CoV-2, indicative of possible conformational differences in the neutralizing epitopes (*9*). More recent studies have reported two SARS-CoV neutralizing antibodies, 47D11 and S309, that have been shown to neutralize SARS-CoV-2 as well (*10, 11*), suggesting that broad cross-neutralizing epitopes exist within lineage B. Convalescent plasma containing SARS-CoV-2 neutralizing antibodies (NAbs) have been shown to confer clear protection in COVID-19 patients (*12, 13*), yet gaps in our knowledge concerning the immunogenic features and key epitopes of SARS-CoV-2, have hampered the development of effective immuno-therapeutics against the virus.

The RBDs of SARS-CoV and SARS-CoV-2 have an amino-acid sequence identity of around 75%, raising the possibility that RBD-targeting cross-neutralizing NAbs could be possibly identified. Using phage display technique, we constructed an antibody library which was generated from RNAs extracted from peripheral lymphocytes of mice immunized with recombinant SARS-CoV RBD. SARS-CoV-2 RBD was used as the target for screening the phage antibody library for potential hits. Antibodies showing tight binding for SARS-CoV-2 RBD were further propagated as chimeric antibodies and tested for neutralizing activities using a vesicular stomatitis virus (VSV) based pseudotyping system (fig. S1) (*14*). Among the antibodies tested, clone 014 which showed potent neutralizing activity against SARS-CoV-2 pseudovirus was humanized, named as H014. To evaluate the binding affinities, real-time association and dissociation of H014 binding to either SARS-CoV-2 RBD or SARS-CoV RBD were monitored using the OCTET system (Fortebio). Both H014 Ig G and Fab fragments exhibited tight binding to both RBDs with comparable binding affinities at sub-nM levels for SARS-CoV-2 RBD and SARS-CoV RBD, respectively (Fig. 1A and fig. S2). Pseudovirus neutralization assays revealed that H014 has potent neutralizing activities: a 50% neutralizing concentration value (IC_50_) of 3 nM and 1 nM against SARS-CoV-2 and SARS-CoV pseudoviruses, respectively (Fig. 1B). Plaque-reduction neutralization test (PRNT) conducted against an authentic SARS-CoV-2 strain (BetaCoV/Beijing/AMMS01/2020) verified the neutralizing activities with an IC_50_ of 38 nM, 10-fold lower than those observed in the pseudotyping system (Fig. 1C). We next sought to assess *in vivo* protection efficacy of H014 in our previously established human ACE2 (hACE2) humanized mouse model that was sensitized to SARS-CoV-2 infection (*15*). In this model, as a result of hACE2 expression on lung cells, SARS-CoV-2 gains entry into the lungs and replicates as in a human disease, exhibiting lung pathology at 5 days post infection (dpi). hACE2-humanized mice were treated by intraperitoneal injection of H014 at 50 mg/kg either 4 hours after (1 dose, therapeutic) or 12 hours before and 4 hours after (2 doses, prophylactic plus therapeutic) intranasal infection with 5 × 10^5^ PFU of SARS-CoV-2 (BetaCoV/Beijing /AMMS01/2020). All challenged animals were sacrificed at day 5. While the viral loads in the lungs of the PBS group (control) surged to ∼10^7^ RNA copies/g at day 5 (Fig. 1D), remarkably, in the prophylactic and prophylactic plus therapeutic groups, H014 treatment resulted in a ∼10-fold and 100-fold reduction of viral titers in the lungs at day 5, respectively (Fig. 1D). Lung pathology analysis showed that SARS-CoV-2 caused mild interstitial pneumonia characterized by inflammatory cell infiltration, alveolar septal thickening and distinctive vascular system injury upon PBS treatment. In contrast, no obvious lesions of alveolar epithelial cells or focal hemorrhage were observed in the lung sections from mice that received H014 treatment (Fig. 1E), indicative of a potential therapeutic role for H014 in curing COVID-19.

**Fig. 1.**
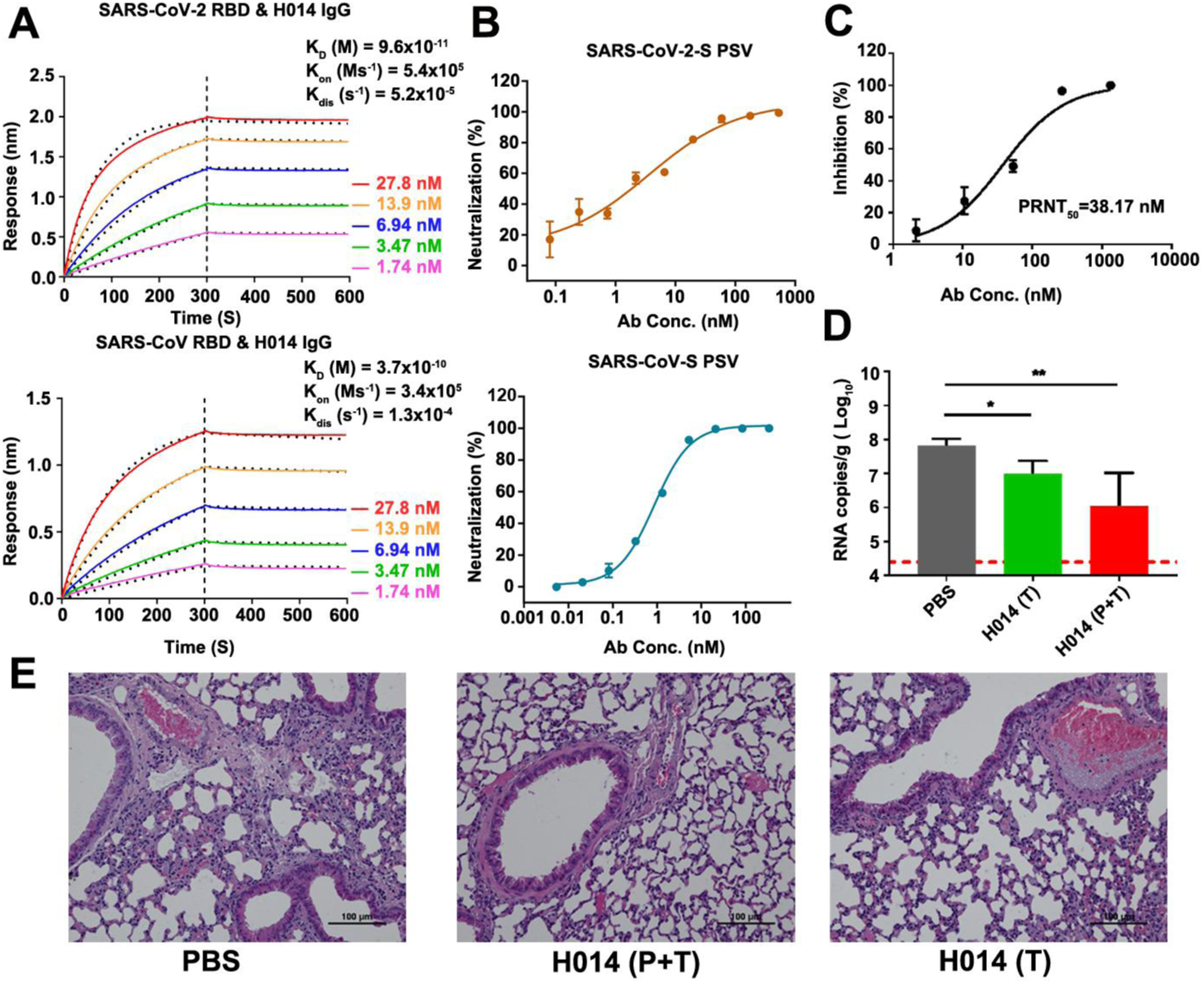
H014 is a lineage B cross-neutralizing antibody of therapeutic value. **(A)** Affinity analysis of the binding of H014 to RBD of SARS-CoV-2 and SARS-CoV. Biotinylated RBD proteins of (upper) SARS-CoV-2 or (lower) SARS-CoV were loaded on Octet SA sensor and tested for real-time association and dissociation of the H014 antibody. Global fit curves are shown as black dotted lines. The vertical dashed lines indicate the transition between association and disassociation phases. **(B)** Neutralizing activity of H014 against SARS-CoV-2 and SARS-CoV pseudoviruses (PSV). Serial dilutions of H014 were added to test its neutralizing activity against (upper) SARS-CoV-2 and (lower) SARS-CoV PSV. Neutralizing activities are represented as mean ± SD. Experiments were performed in triplicate. **(C)** *In vitro* neutralization activity of H014 against SARS-CoV-2 by PRNT in Vero cells. Neutralizing activities are represented as mean ± SD. Experiments were performed in duplicates. **(D)** Groups of hACE2 mice that received SARS-CoV-2 challenge were treated intraperitoneally with H014 in two independent experimental settings: 1) a single dose at 4 h post infection (Therapeutic, T); 2) two doses at 12 h before and 4 h post challenge (Prophylactic plus Therapeutic, P+T). Virus titers in the lungs were measured 5 days post infection (dpi) and are presented as RNA copies per gram of lung tissue. n=7/3/3, respectively. *P<0.05. LOD represents limit of detection. **(E)** Histopathological analysis of lung samples at 5 dpi. Scale bar: 100 µm.

The overall structure of SARS-CoV-2 S trimer resembles those of SARS-CoV and other coronaviruses. Each monomer of the S protein is composed of two functional subunits. The S1 subunit binds the host cell receptor, while the S2 subunit mediates fusion of the viral membrane with the host cell membrane (*2, 16*). The four domains within S1 include N-terminal domain (NTD), RBD and two subdomains (SD1 and SD2) where the latter are positioned adjacent to the S1/S2 cleavage site. Hinge-like movements of the RBD give rise to two distinct conformational states referred to as the “close” and “open”, where close corresponds to the receptor-inaccessible state and open corresponds to the receptor-accessible state, which is supposed to be metastable (*17-20*). Cryo-EM characterization of the stabilized SARS-CoV-2 S ectodomain in complex with the H014 Fab fragment revealed that the complex adopts three distinct conformational states, corresponding to one RBD open + two RBDs closed (state 1), two RBDs open + one RBD closed (state 2) and all three open RBDs (state 3) (Fig. 2A). Interestingly, structure of the completely closed (state 4) SARS-CoV-2 S trimer without any Fab bound was also observed during 3D classification of the cryo-EM data, albeit in the presence of excessive Fab, suggesting that the binding sites of H014 are exposed through protein “breathing” followed by a stochastic RBD movement (Fig. 2A). We determined asymmetric cryo-EM reconstructions of the 4 states at 3.4 – 3.6 Å (figs. S3-S6 and table S1). However, the electron potential maps for binding interface between RBD and H014 are relatively weak due to conformational heterogeneity. To solve this problem, focusing classification and refinement by using a “block-based” reconstruction approach were performed to further improve the local resolution up to 3.9 Å, enabling reliable analysis of the interaction mode (fig. S5-S6). Detailed analysis of the interactions between H014 and S was done using the binding interface structure.

**Fig. 2.**
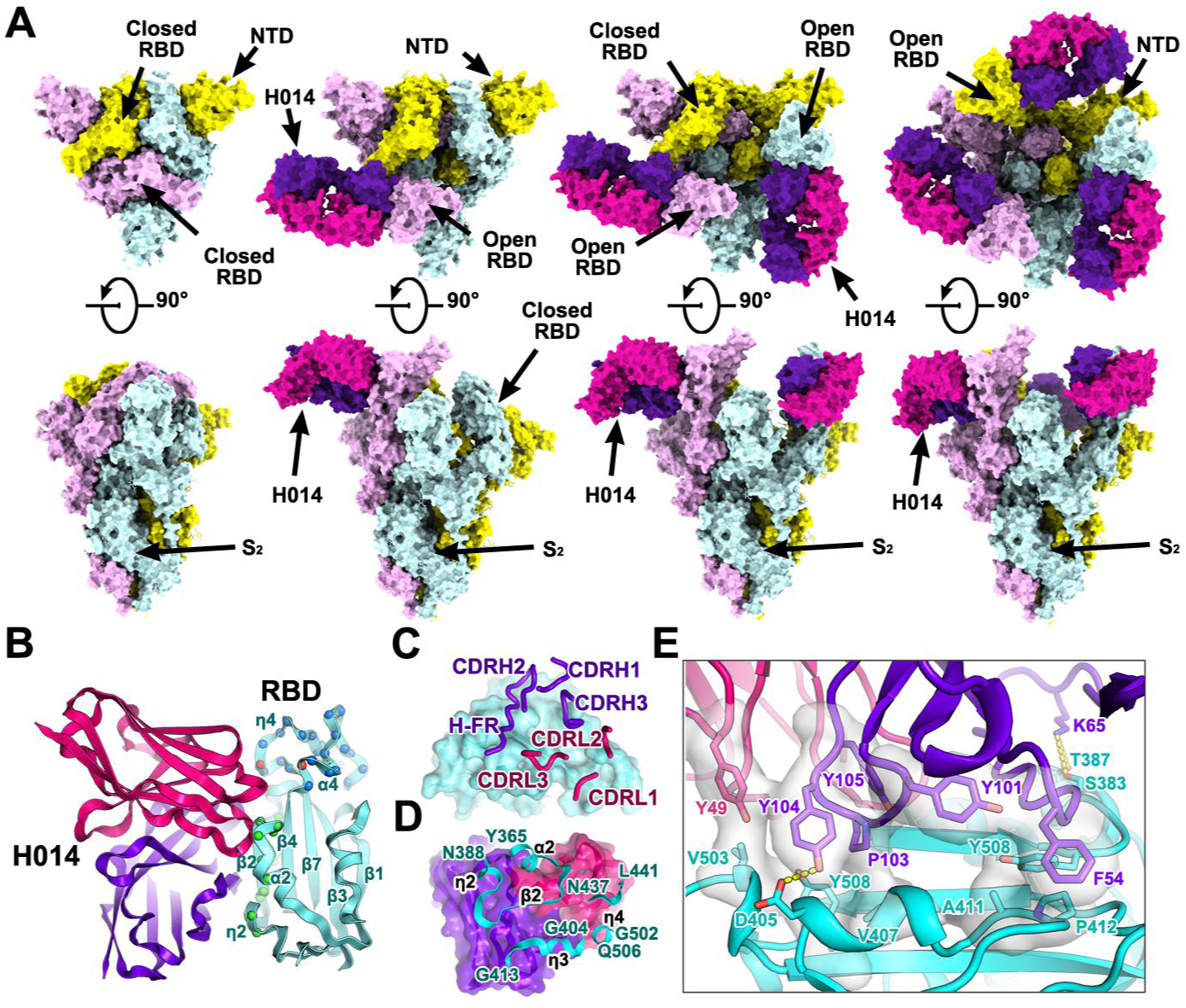
Cryo-EM structures of the SARS-CoV-2 S trimer in complex with H014. **(A)** Orthogonal views of SARS-CoV-2 S trimer with three RBDs in the closed state (left), one RBD in the open state and complexed with one H014 Fab (middle), two RBDs in the open state and each complexed with one H014 Fab. NTD: N-terminal domain. All structures are presented as molecular surfaces with different colors for each S monomer (cyan, violet and yellow), and the H014 Fab light (hotpink) and heavy (purpleblue) chains. **(B)** Cartoon representations of the structure of SARS-CoV-2 RBD in complex with H014 Fab with the same color scheme as in Fig. 2A. Residues comprising the H014 epitope and the RBM are shown as spheres and colored in green and blue, respectively. The overlapped residues between the H014 epitope and the RBM are shown in red. **(C)** and **(D)** Interactions between the H014 and SARS-CoV-2 RBD. The CDRs of the H014 that interact with SARS-CoV-2 RBD are displayed as thick tubes over the cyan surface of the RBD **(C)**. The H014 epitope is shown as a cartoon representation over the surface of the RBD **(D). (E)** Details of the interactions between the H014 and SARS-CoV-2 RBD. Some residues involved in the formation of hydrophobic patches and hydrogen bonds are shown as sticks and labeled. Color scheme is the same as in Fig. 2A.

H014 recognizes a conformational epitope on one side of the open RBD, only involving protein/protein contacts, distinct from the receptor-binding motif (RBM) (Fig. 2B). The H014 paratope constitutes all six complementary determining region (CDR) loops (CDRL1-3 and CDRH1-3) and unusual the heavy-chain frame work (HF-R, residues 58-65) that forge tight interactions with the RBD, resulting in a buried area of ∼1,000 Å^2^ (Fig. 2C). Variable domains of the light-chain and heavy-chain contribute ∼32% and 68% of the buried surface area, respectively, through hydrophobic and hydrophilic contacts. The H014 epitope is composed of 21 residues, primarily locating in the α2-β2-η2 (residues 368-386), η3 (residues 405-408, 411-413), α4 (residue 439) and η4 (residues 503) regions, which construct a cavity on one side of the RBD (Fig. 2B, 2D and fig. S7). The 12-residue long CDRH3 inserts into this cavity and the hydrophobic residues (YDPYYVM) enriched CDRH3 contacts the η3 and edge of the five-stranded β-sheet (β 2) region of the RBD (Fig. 2D). Tight bindings between the RBD and H014 are primarily due to extensive hydrophobic interactions contributed by two patches: one formed by F54 from CDRH2, Y101 from CDRH3 and A411, P412 and Y508 of the RBD and the other composed of Y49 from CDRL2, P103, Y104, Y105 from CDRH3 and V407, V503 and Y508 of the RBD (Fig. 2E and table S2). Additionally, hydrophilic contacts from CDRH1 and HF-R further enhance the RBD-H014 interactions, leading to an extremely high binding affinity at sub nM level at temperatures of 25 °C or 37 °C (Fig. 2E and fig. S8). Residues comprising the epitope are mostly conserved with 3 single-site mutants (R408I, N439K and V503F) in this region among currently circulating SARS-CoV-2 strains reported (fig. S9). In addition, a number of SARS-CoV-2 isolates bear a common mutation V367F in the RBD (*21*), which lies adjacent to the major epitope patch α 2-β2-η2. Constructs of the recombinant RBD harboring point mutations of above residues and other reported substitutions exhibited an indistinguishable binding affinity to H014 (fig. S10), suggesting that H014 may exhibit broad neutralization activities against SARS-CoV-2 strains currently circulating worldwide. Out of the 21 residues in the H014 epitope, 17 (81 %) are identical between SARS-CoV-2 and SARS-CoV (fig. S9), which explains the cross-reactivity and comparable binding affinities.

To investigate whether H014 interferes with the binding of RBDs of SARS-CoV-2 or SARS-CoV to ACE2, we performed competitive binding assays at both protein and cellular levels. The enzyme-linked immunosorbent assay (ELISA) indicated that H014 was able to compete with recombinant ACE2 for binding to RBDs of SARS-CoV-2 and SARS-CoV with EC_50_ values of 0.7 nM and 5 nM, respectively (Fig. 3A). Additionally, H014 efficiently blocked both the attachment of SARS-CoV-2 RBD to ACE2 expressing 293T cells and the binding of recombinant ACE2 to SARS-CoV-2 S expressing 293T cells (Fig. 3B). To verify its potential full occlusion on trimeric S, we conducted two sets of surface plasmon resonance (SPR) assays by exposing the trimeric S to H014 first and then to ACE2 or the other way around. As expected, binding of H014 completely blocked the attachment of ACE2 to trimeric S. Moreover, ACE2 could be displaced from trimeric S and replaced by H014 (Fig. 3C). To further verify these results in a cell-based viral infection model, real time RT-PCR analysis was carried out to quantify the amount of virus remaining on the host cell surface, which were exposed to antibodies pre- or post-virus attachment to cells at 4 °C. H014 efficiently prevented attachment of SARS-CoV-2 to the cell surface in a dose-dependent manner and the viral particles that had already bound to the cell surface could be partially stripped by H014 (Fig. 3D). Superimposition of the structure of the H014-SARS-CoV-2 trimeric S complex over the ACE2-SARS-CoV-2 RBD complex structure revealed clashes between the ACE2 and H014, arising out of an overlap of the regions belonging to the binding sites located at the apical helix (η4) of the RBD (Fig. 3E). This observation differs substantially from those of most known SARS RBD-targeted antibody complexes, where the antibodies directly recognize the RBM (*22-25*). Thus, the ability of H014 to prevent SARS-CoV-2 from attaching to host cells can be attributed to steric clashes with ACE2.

**Fig. 3.**
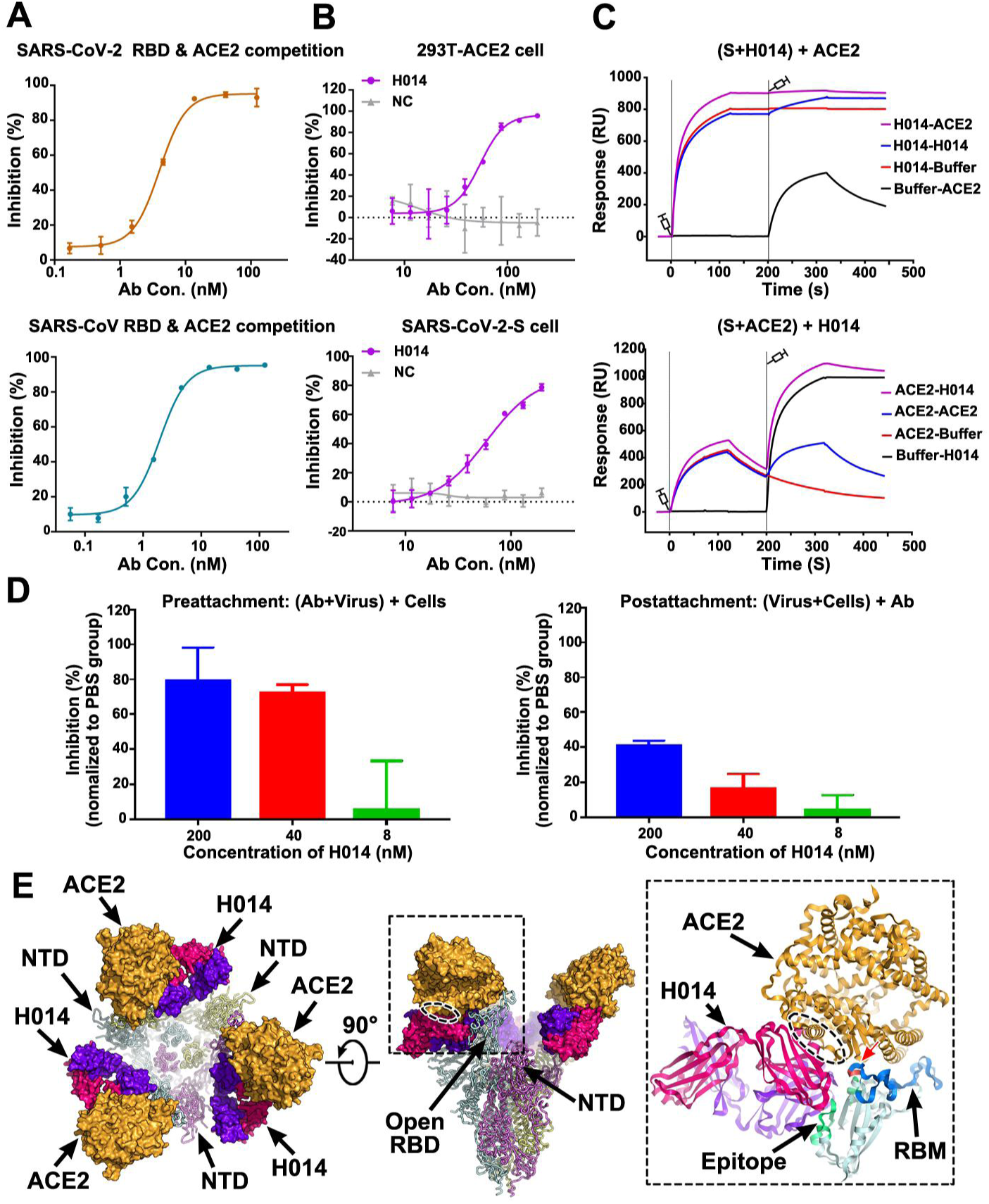
Mechanism of neutralization of H014. **(A)** Competitive binding assays by ELISA. Recombinant SARS-CoV-2 (upper) or SARS-CoV (lower) RBD protein was coated on 96-well plates, recombinant ACE2 and serial dilutions of H014 were then added for competitive binding to SARS-CoV-2 or SARS-CoV RBD. Values are mean ± SD. Experiments were performed in triplicate. **(B)** Blocking of SARS-CoV-2 RBD binding to 293T-ACE2 cells by H014 (upper). Recombinant SARS-CoV-2 RBD protein and serially diluted H014 were incubated with ACE2 expressing 293T cells (293T-ACE2) and tested for binding of H014 to 293T-ACE2 cells. Competitive binding of H014 and ACE2 to SARS-CoV-2-S cells (lower). Recombinant ACE2 and serially diluted H014 were incubated with 293T cells expressing SARS-CoV-2 Spike protein (SARS-CoV-2-S) and tested for binding of H014 to SARS-CoV-2-S cells. BSA was used as a negative control (NC). Values are mean ± SD. Experiments were performed in triplicate. **(C)** BIAcore SPR kinetics of competitive binding of H014 and ACE2 to SARS-CoV-2 S trimer. For both panels, SARS-CoV-2 S trimer was loaded onto the sensor. In the upper panel, H014 was first injected, followed by ACE2, whereas in the lower panel, ACE2 was injected first and then H014. The control groups are depicted by black curves. **(D)** Amount of virus on the cell surface, as detected by RT-PCR. Pre-attachment mode: incubate SARS-CoV-2 and H014 first, then add the mixture into cells (left); post-attachment mode: incubate SARS-CoV-2 and cells first, then add H014 into virus-cell mixtures (right). High concentrations of H014 prevent attachment of SARS-CoV-2 to the cell surface when SARS-CoV-2 was exposed to H014 before cell attachment. Values represent mean ± SD. Experiments were performed in duplicates. **(E)** Clashes between H014 Fab and ACE2 upon binding to SARS-CoV-2 S. H014 and ACE2 are represented as surface; SARS-CoV-2 S trimer is shown as ribbon. Inset is a zoomed-in view of the interactions of the RBD, H014 and ACE2 and the clashed region (oval ellipse) between H014 and ACE2. The H014 Fab light and heavy chains, ACE2, and RBD are presented as cartoons. The epitope, RBM and the overlapped binding region of ACE2 and H014 on RBD are highlighted in green, blue and red, respectively.

Similar to the RBM, the H014 epitope is only accessible in the open state, indicative of a role akin to the RBM *–* involving dynamic interferences in interactions with host cells. Our structures together with previously reported coronavirus S structures, not including human coronavirus HKU1 (*20*), have observed the breathing of the S1 subunit, that mediates the transition between “close” and “open” conformation (Fig. 4A) (*2, 9, 20*). In contrast to the hinge-like movement of the RBD observed in most structures, the conformational transition from “close” to “open” observed in our structures mainly involves two steps of rotations: 1) counterclockwise movement of SD1 by ∼25° encircling the hinge point (at residue 320); 2) the RBD rotates itself counterclockwise by ∼60° (Fig. 4B). These conformational rotations of the SD1 and RBD at proximal points relay an amplified alteration at the distal end, leading to opening up of adequate space for the binding of H014 or ACE2 (Fig. 4B). Ambiguously, the special conformational transition observed in our complex structures results from the engagement of H014 or a synergistic movement of the SD1 and RBD.

**Fig. 4.**
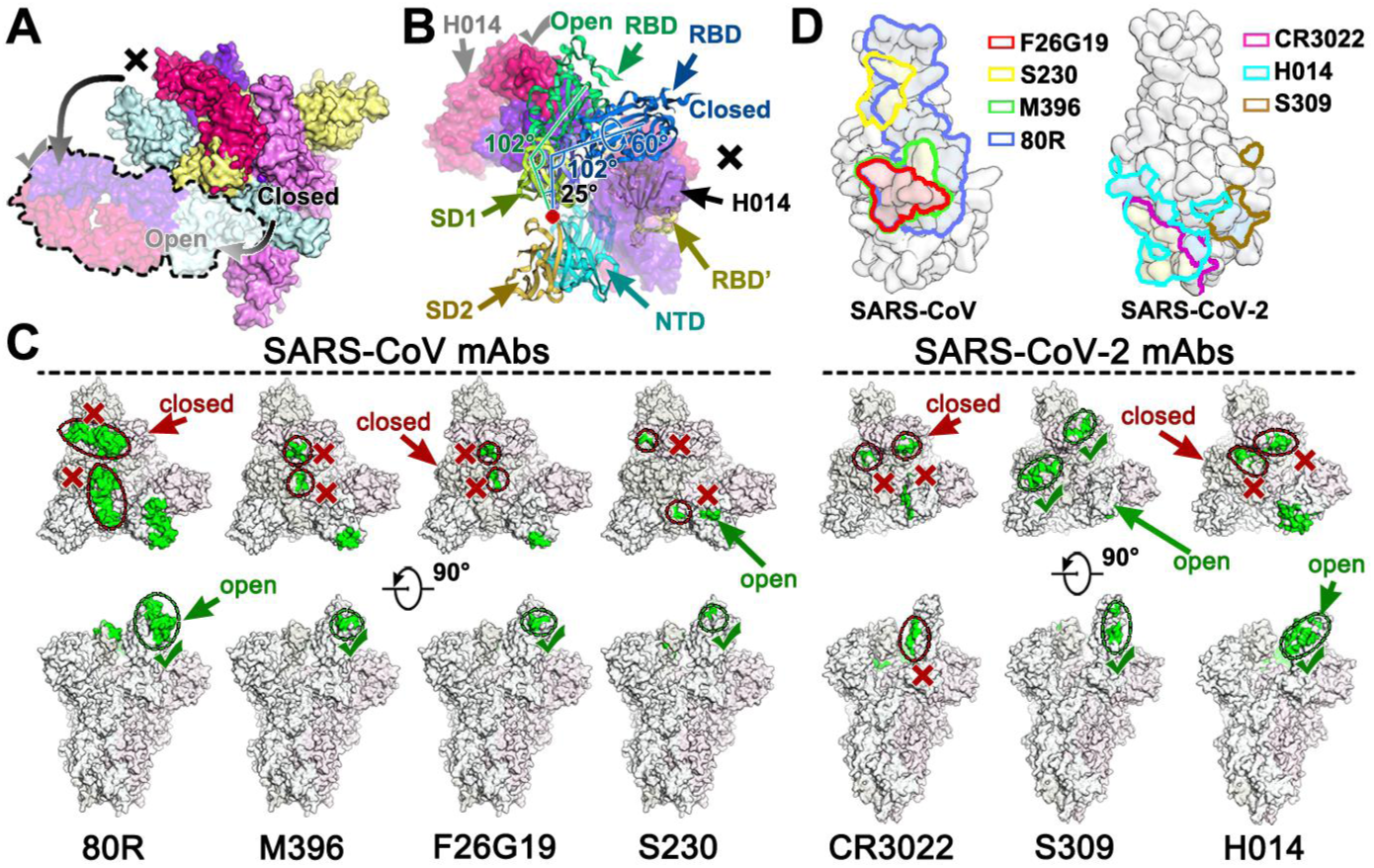
Breathing of the S1 subunit and epitopes of neutralizing antibodies. **(A)** H014 can only interact with the “open” RBD, whereas the “closed” RBD is inaccessible to H014. The “open” RBD and RBD bound H014 are depicted in lighter colors corresponding to the protein chain they belong to. Color scheme is the same as in Fig. 2A. **(B)** Structural rearrangements of the S1 subunit of SARS-CoV-2 transition from the closed state to the open state. SD1: subdomain 1, SD2: subdomain 2, RBD’: RBD (closed state) from adjacent monomer. SD1, SD2, NTD, RBD and RBD’ are colored in pale green, light orange, cyan, blue and yellow, respectively. The red dot indicates the hinge point. The angles between the RBD and SD1 are labeled. **(C)** Epitope location analysis of neutralizing antibodies on SARS-CoV and SARS-CoV-2 S trimers. The S trimer structures with one RBD open and two RBD closed from SARS-CoV and SARS-CoV-2 were used to show individual epitope information, which is highlighted in green. The accessible and in-accessible states are encircled and marked by green ticks and red crosses. **(D)** Footprints of the seven mAbs on RBDs of SARS-CoV (left) and SARS-CoV-2 (right). RBDs are rendered as molecular surfaces in light blue. Footprints of different mAbs are highlighted in different colors as labeled in the graph. Note: epitopes recognized by indicated antibodies are labeled in blue (non-overlapped), yellow (overlapped once) and red (overlapped twice).

Humoral immunity is essential for protection against coronavirus infections and passive immunization of convalescent plasma has been demonstrated to be effective in treating SARS and COVID-19 (*12, 26*). A number of SARS-CoV specific neutralizing antibodies including m396, 80R and F26G19, have been previously reported to engage primarily the RBM core (residues 484-492 in SARS-CoV) provided that the RBD is in open conformation (Fig. 4C and 4D) (*24, 27, 28*). Unexpectedly, none of these antibodies has so far demonstrated impressive neutralizing activity against SARS-CoV-2 (*7, 8*). S230, a SARS-CoV neutralizing antibody that functionally mimics receptor attachment and promotes membrane fusion, despite requiring an open state of the RBD as well, recognizes a small patch at the top of the RBM (Fig. 4C and 4D) (*17*). Recently, two other antibodies (CR3022 and S309) originating from the memory B cells of SARS survivors in 2003, demonstrated strong binding to SARS-CoV-2 (*9, 10*). Surprisingly, CR3022 failed to neutralize SARS-CoV-2 *in vitro*, which could be attributed to the fact that CR3022 binds a cryptic epitope, which is distal from the RBD and is only accessible when at least two RBDs are in the open state. Furthermore, CR3022 does not compete with ACE2 for binding to the RBD (Fig. 4C and 4D) (*9*). By contrast, S309, a more recently reported neutralizing mAb against both SARS-CoV and SARS-CoV-2, recognizes a conserved glycan-containing epitope accessible in both the open and closed states of SARS-CoV and SARS-CoV-2 RBDs (*10*). The neutralization mechanism of S309 does not depend on direct blocking of receptor binding, but it induces antibody-dependent cell cytotoxicity (ADCC) and antibody-dependent cellular phagocytosis (ADCP). We reported a potent therapeutic antibody H014, which can cross-neutralize SARS-CoV-2 and SARS-CoV infections by blocking attachment of the virus to the host cell (Fig. 3). Distinct from other antibodies, H014 binds one full side of the RBD, spanning from the proximal η 2 to the distal edge of the RBM and possesses partially overlapping epitopes with SARS-CoV specific antibodies like m392, 80R, F26G19, CR3022 and S309 (Fig. 4C and 4D). Although both SARS-CoV and SARS-CoV-2 utilize ACE2 to enter host cells, high sequence variations in the RBM and local conformational rearrangements at the distal end of the RBM from the two viruses have been observed (*8*). These probably lead to the failure of some antibodies, especially those targeting the RBM of SARS-CoV, in cross-binding and cross-neutralizing SARS-CoV-2. Three cross-reactive antibodies (CR3022, S309 and H014) recognize more conserved epitopes located beyond the RBM, amongst which the CR3022 epitope is most distant from the RBM (Fig. 4C and 4D), suggesting a possible reason for its inability to neutralize SARS-CoV-2. H014 binds more conserved epitopes, some of which are in close proximity to the RBM, rendering it effective in cross-neutralizing lineage B viruses. Additionally, antibodies recognizing more conserved patches beyond the RBM would function synergistically with antibodies targeting the RBM to enhance neutralization activities, which could mitigate the risk of viral escape. The molecular features of H014 epitopes unveiled in this study facilitate the discovery of broad cross-neutralizing epitopes within lineage B and pose interesting targets for structure-based rational vaccine design. Our studies also highlight the promise of antibody-based therapeutic interventions for the treatment of COVID-19.

## Acknowledgments

We thank Dr. Xiaojun Huang, Dr. Boling Zhu and Dr. Gang Ji for cryo-EM data collection, the Center for Biological imaging (CBI) in Institute of Biophysics for EM work. Work was supported by the National Key Research and Development Program (2020YFA0707500, 2018YFA0900801), the Strategic Priority Research Program (XDB29010000), National Science and Technology Major Projects of Infectious Disease funds (grants 2017ZX103304402) and Beijing Municipal Science and Technology Project (Z201100005420017). Xiangxi Wang was supported by Ten Thousand Talent Program and the NSFS Innovative Research Group (No. 81921005). Cheng-Feng Qin was supported by the National Science Fund for Distinguished Young Scholar (No. 81925025) and the Innovative Research Group (No. 81621005) from the NSFC, and the Innovation Fund for Medical Sciences (No.2019-I2M-5-049) from the Chinese Academy of Medical Sciences. ” L.X. and C. S. are inventors on patent application (202010219867.1) submitted by Sinocelltech. Ltd that covers the intellectual property of H014”.

## Author contributions

Zhe.L., Y.D., C.S., Q.Y., L.C., C.F., W.H., Y.S., Q.C., S.S., N.W., D.Z., J.N., Z.C., N.S. and X.L. performed experiments; X.W., C-F.Q, Z.R., L.X. and Y.W. designed the study; all authors analyzed data; and X.W., L.Z., C-F.Q. and Z.R. wrote the manuscript.

## Data and materials availability

Cryo-EM density maps of the *apo* SARS-CoV-2 S trimer, SARS-CoV-2 S trimer in complex with one Fab, SARS-CoV-2 S trimer in complex with two Fabs, SARS-CoV-2 S trimer in complex with three Fabs and binding interface have been deposited at the Electron Microscopy Data Bank with accession codes EMD-30325, EMD-30326, EMD-30332, EMD-30333 and EMD-30331 and related atomic models has been deposited in the protein data bank under accession code 7CAB, 7CAC, 7CAI, 7CAK and 7CAH, respectively.

