## supplementary materials for "Structural basis for neutralization of SARS-CoV-2 and SARS-CoV by a potent therapeutic antibody"

**This PDF file includes:**

Materials and Methods

Figs. S1 to S10

Tables. S1 to S2

### Materials and Methods

#### Facility and ethics statements

All experiments with live SARS-CoV-2 viruses were performed in the enhanced biosafety level 3 (P3+) facilities of the Institute of Microbiology and Epidemiology, Academy of Military Medical Sciences. All animal experiments were approved by the Experimental Animal Committee of Laboratory Animal Center, AMMS (approval number: IACUC-DWZX-2020-001).

#### Cells and viruses

Vero cells (ATCC, CCL-81) were maintained in Dulbecco's Modified Essential Medium supplemented with 10% fetal bovine serum (FBS) (Biowest) at 37 °C with 5% $CO_2$. The strain of BetaCoV/wuhan/AMMS01/2020 was originally isolated from a COVID-19 patient returning from Wuhan, China. The virus was amplified and titrated by standard plaque forming assay on Vero cells.

#### Protein expression and purification

The plasmids used for protein expression were individually constructed by insertion of the coding sequences for SARS-CoV RBD (residues 306–527, GenBank: NC_004718.3), SARS-CoV-2 RBD (residues 319-541, GenBank: MN908947.3), and SARS-CoV-2 S trimer (residues 1–1208, GenBank:MN908947.3) into the mammalian expression vector pCAGGS with a C-terminal 2×StrepTag to facilitate protein purification. The gene of S protein was constructed with proline substitutions at residues 986 and 987, a "GSAS" instead of "RRAR at the furin cleavage site according to Jason S. McLellan's research (*16*). SARS-CoV RBD, SARS-CoV-2 RBD and SARS-CoV-2 S trimer were used to transiently transfect HEK Expi 293F cells (Thermo Fisher) using polyethylenimine. Protein was purified from filtered cell supernatants using StrepTactin resin (IBA) before being subjected to additional purification by size-exclusion chromatography using a Superose 6 10/300 column (GE Healthcare) or a Superdex 200 10/300 Increase column (GE Healthcare) in 20mM Tris, 200 mM NaCl, pH 8.0.

#### Generation of humanized anti-SARS-CoV-2 antibody H014

Antibodies against SARS-CoV-2 RBD were screened from a phage-display scFv library generated from spleen mRNA of mice immunized with recombinant

SARS-CoV RBD. The recombinant SARS-CoV-2 RBD was used as the target protein to select specific anti-RBD scFvs via biopanning. Candidate scFvs with potent binding to recombinant SARS-CoV-2 RBD were further produced as chimeric antibodies. The chimeric mh014 showed high affinity for RBDs of both viruses and potent neutralizing activity against both pseudoviruses. The antibody was further humanized to generate humanized antibody H014 (IgG1 subtype).

**Protein-protein interactions identified by Octet**

Recombinant SARS-CoV-2 RBD or SARS-CoV RBD was biotinylated and loaded using SA sensor (Pall corporation). H014 were then added for real-time association and dissociation analysis using Octet96e at room temperature (Fortebio). Data Analysis Octet was used for data processing.

**Pseudovirus production in 293T cells**

Pseudoviruses were generated as previously described (*14*). Briefly, 293T cells were transfected with the plasmid of SARS-CoV-2 S or SARS-CoV S, respectively. 24 h later, transfected 293T cells were infected with VSV G pseudotyped virus (G*ΔG-VSV) at a multiplicity of infection (MOI) of 4. Two hours after infection, cells were washed with PBS three times, and then complete culture medium was added. Twenty-four hours post infection, SARS-CoV-2 or SARS-CoV pseudovirus was harvested, clarified by passing through 0.45-μm filters and stored at −80°C until further use.

**Pseudovirus neutralization assay**

Aliquots of 100μL of ~40,000 Vero cells/well were added into a 96-well plate. PSV-mAb complexes were obtained by incubating 60 μL of SARS-CoV-2/ SARS-CoV PSV with 60 μL of serial dilutions of antibody samples for 1 h at 37 °C. Then the antibody-PSV mixtures were added to Vero cells. Following 24h of incubation in a 5% $CO_2$ environment at 37 °C the luciferase luminescence (RLU) was measured using luciferase assay system according to the manufacturer's procedure with a luminescence microplate reader. The neutralization percentage was calculated as following: Inhibition (%) = [1- (sample RLU- Blank RLU) / (Positive Control RLU-Blank RLU)] (%). Antibody neutralization titers were presented as 50% maximal inhibitory concentration ($IC_{50}$).

**Flow cytometry**

Indicated concentrations of H014 were incubated with 293T-ACE2 or 293T-SARS-CoV-2-S cells along with recombinant SARS-CoV-2 RBD or ACE2 (Cat:10108-H08H, Sino Biological) for 45 min, respectively. After washing away the unbound proteins, cells were incubated with FITC labeled secondary antibody for 20 mins and passed through flow cytometer for detection of cellular binding. Flowjo and Graphpad were used for data analysis.

**Production of Fab fragment**

The H014 Fab fragment was generated using a Pierce FAB preparation Kit (Thermo Scientific), according to the manufacturer's instructions. Briefly, after removal of the salt using a desalting column, the antibody was mixed with papain and then digested at 37 ˚C for 3-4 h. The Fab was separated from the Fc fragment by protein A affinity column and then concentrated for cryo-EM analysis.

**Negative stain**

Samples to be examined were diluted to an appropriate concentration (~0.02 mg/mL) and dropped onto a freshly glow-discharged carbon-coated grid. After rinsing twice with buffer (20 mM Tris, 200 mM NaCl, pH 8.0), the grid was stained with 1% phosphotungstic acid (pH 7.0) and then loaded onto a 120 kV TEM for examination.

**Cryo-EM sample preparation and data collection**

3 μL of purified SARS-CoV-2 S at 1 mg/ml was mixed with 2 μL of H014 Fab fragments at 1.1 mg/ml in purification buffer solution (20 mM Tris, 200 mM NaCl, pH 8.0) and incubated for 2 min at room temperature. A 3 μL aliquot of the mixture was transferred onto a freshly glow-discharged C-flat R1.2/1.3 Cu grid. Grids were blotted for 3 s in 100% relative humidity for plunge-freezing (Vitrobot; FEI) in liquid ethane. Cryo-EM data sets were collected at 300 kV using a Titan Krios microscope (Thermo Fisher) equipped with a K2 detector (Gatan, Pleasanton,CA). Movies (32 frames, each 0.2 s, total dose 60 $e^{-}Å^{-2}$) were recorded with a defocus of between 1.5 and 2.7 μm using SerialEM, which yields a final pixel size of 1.04 Å.

**Image processing**

A total of 12,146 micrographs were recorded for SARS-CoV-2 S trimer-H014-Fab complex. Out of these, 11,504 micrographs with visible CTF rings beyond 1/5 Å in their spectra were selected for further processing. The defocus value for each micrograph was determined using Gctf (*29*). Then particles were picked and extracted for two-dimensional alignment. The well-defined particle images were selected for three-dimensional reconstruction in Relion (*30*). Subsequently, apart from the rubbish class (~9% particles), four major classes: the *apo* S trimer, S trimer with 1 or 2 or 3 Fabs bound (states 1-4) were separated after the 3D classification without imposing symmetry, using previously reported closed SARS-CoV-2 structure (*2*) as initial model. To further separate and improve the reconstruction resolution of the complex structures, a second cycle of 3D classification was carried out. After the high-resolution refinement and postprocessing (estimate the B-factor automatically), the final resolution was evaluated on the basis of the gold-standard Fourier shell correlation (threshold = 0.143) (*31*). Although the overall resolution for these structures (states 1-4) are up to 3.5 Å – 3.6 Å, the maps for the binding interface between RBD and H014 are quite weak due to the conformational heterogeneity of the RBD, which is similar to previous structural investigations. To improve the resolution for the binding interface, we used the block-based reconstruction strategy (*32-34*) for focusing classification and refinement. The orientation parameters of each particle image (states 1-3) determined in Relion were used to guide extraction of the block region (~50% bigger than NTD-RBD-Fab) and these blocks were further classified. A local reconstruction focusing on the NTD-RBD-Fab region was carried out. Furthermore, the density map for the binding interface could be improved further by local averaging of the RBD-Fab equivalent copies present in different classes, finally yielding a resolution of 3.9 Å for the interface. The local resolution was evaluated by ResMap (*35*).

**Model building and refinement**

The structures of the *apo* SARS-CoV-2 S trimer and a human Fab fragment (Protein Data Bank ID: 6VSB and 5N4J) were manually fitted into the refined map of SARS-CoV-2 S trimer-H014 complex using Chimera (*36*) and further corrected manually by real-space refinement in COOT (*37*). The atomic model was further refined by positional and B-factor refinement in real space using Phenix (*38*). The final models were evaluated by Molprobity (*39*). Details of the data sets and refinement statistics are summarized in table S1.

**Surface plasmon resonance**

SARS-CoV-2 S trimer was immobilized onto a CM5 sensor chip surface using the NHS/EDC method to a level of ~600 response units (RUs) using a Biacore 8K (GE Healthcare) and a PBS running buffer (supplemented with 0.05% Tween-20). Serial dilutions of purified H014 were injected in concentration from 125 to 7.8 nM. Serial dilutions of purified and His-tagged ACE2 were injected in concentration from 500 to 31.25 nM. For the competitive binding assays, the first sample flew over the chip at a rate of 20 μl/min for 120 s, then the second sample was injected at the same rate for another 120 s. The response units were recorded at room temperature and analyzed using the same software as mentioned above.

**Plaque reduction neutralization tests (PRNT)**

Standard plaque reduction neutralization tests (PRNT) against SARS-CoV-2 were performed to test the neutralization activity of H014-25 in Vero cells. Briefly, 5-fold serial dilutions of mAbs were added to approximately 100 PFU of SARS-CoV-2 and incubated at 37 °C for 1 h. Then, the mixture was added to Vero cell monolayers in a 12-well plate in duplicate and incubated at 37 °C for 1 h. The mixture was removed, and 1 ml of 1.0% (w/v) LMP agarose (Promega) in DMEM plus 4% (v/v) FBS was layered onto the infected cells. After further incubation at 37 °C for 2 days, the wells were stained with 1% (w/v) crystal violet dissolved in 4% (v/v) formaldehyde to visualize the plaques. The PRNT50 values were determined using non-linear regression analysis using GraphPad prism.

**Evaluation of protection against SARS-CoV-2 by H014 in hACE2 mice model**

The *in vivo* protection efficacy of H014 was assessed by using a humanized hACE2 mouse model (*15*). Briefly, a group of 6-8 week hACE2 humanized mice were intraperitoneally administrated with H014 (50 mg/kg) before (prophylactic) and/or after (therapeutic) challenge with $5 \times 10^5$ PFU of SARS-CoV-2 via intranasal route. All mice were monitored daily for morbidity and mortality. The lung tissues of mice were collected at 5 dpi for viral RNA loads assay and HE staining.

**Viral RNA quantitation**

Viral RNA quantification was performed by RT-qPCR using One Step PrimeScript RT-PCR Kit (Takara, Japan). The primers and probe targeting the spike (S) gene of SARS - CoV - 2 used for RT-qPCR were CoV-F3 (5‘-TCCTGGTGATTCTT CTTCAGGT-3’); CoV-R3 (5‘-TCTGAGAGAGGGTCAAGTGC-3’); and CoV-P3 (5‘-AGCTGCAGCACCAGCTGTCCA-3’), respectively.

**RT-PCR quantification of the virus on the cell surface**

Pre- and post-adsorption inhibition assay were performed as described previously (40). In the post-adsorption assay, SARS-CoV-2 firstly was added to Vero cells for 1 h at 4 °C. Then the cells were washed three times. After this step the mAb was added and the cells were incubated for an additional 1 h at 4 °C. In the pre-adsorption assay, the mAb was incubated with Vero cells for 1 h at 4 °C after which SARS-CoV-2 was added. After three washes with PBS, the PRNT assay was performed as described above. Quantitative RT-PCR was performed to detect the amount of RNA of SARS-CoV-2 remaining on the surface of Vero cells after H014 treatment.

**Histology and Immunostaining**

Mouse tissues were excised and fixed with 10% neutral buffered formaline, dehydrated and embedded in paraffin. Sections at a thickness of 4 μm were stained with hematoxylin and eosin (H & E) according to standard histological procedures. Images were captured using Olympus BX51 microscope equipped with a DP72 camera.

**Binding of H014 to SARS-CoV-2 RBD mutants by ELISA**

Various SARS-CoV-2 RBD recombinant proteins harboring previously reported point mutations were generated and tested for H014 binding by ELISA. Briefly, RBD mutant proteins were coated on 96-well plates using PBS buffer over night at 4 °C. BSA was used for blocking at room temperature for 1 h. Serial dilutions of antibodies were then added and incubated at room temperature for 1 h. After washing away the unbound antibodies, secondary antibody against human Fab with HRP label were added and incubated for another one hour before washing away the excess labelled antibody. For color development, TMB solution was added and incubated for 5-30 min followed by addition of 1% $H_2SO_4$ to stop the reaction. Absorbance at 450 nm was monitored using a microplate reader.

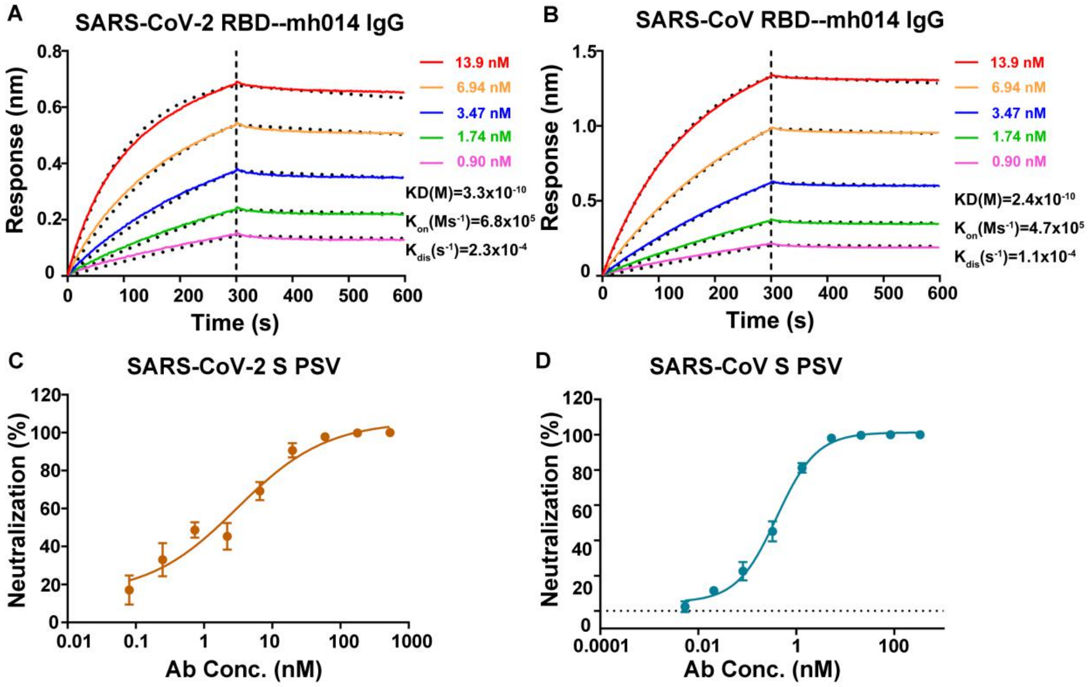


**Fig S1. Cross binding and neutralization activity of mh014 against SARS-CoV-2 and SARS-CoV.**

**(A-B)** Binding affinities of mh014 to the RBD of SARS-CoV-2 and SARS-CoV by Octet system. Biotinylated RBD proteins of SARS-CoV-2 (**A**) or SARS-CoV (**B**) were loaded on Octet SA sensor and detected by real-time association and dissociation of H014 antibody. Global fit curves are shown as black dotted lines. The vertical dashed lines indicate the transition between association and disassociation phases. **(C-D)** Neutralizing activities of mh014 against SARS-CoV-2 and SARS-CoV pseudoviruses (PSV). Serial dilutions of mh014 were added to test its neutralizing activity against SARS-CoV-2 PSV (**C**) and SARS-CoV PSV (**D**).

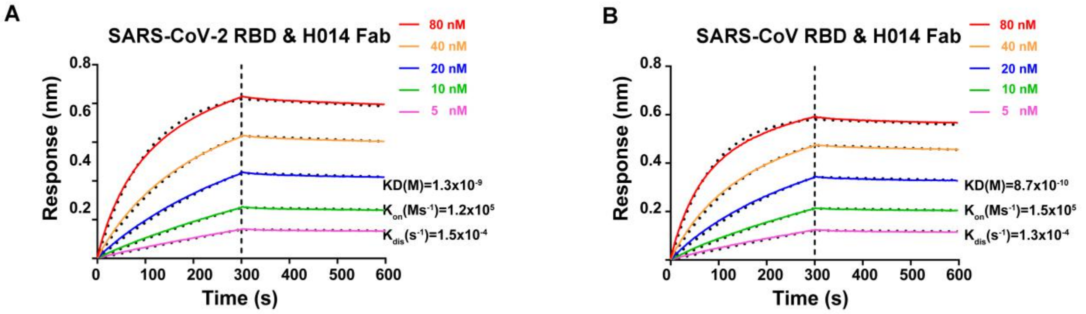


**Fig S2. Binding affinities of H014 Fab fragments to SARS-CoV-2 and SARS-CoV**

Affinity analysis of the binding of H014 Fab fragments to SARS-CoV-2 RBD **(A)** and SARS-CoV RBD **(B)**. Biotinylated RBD proteins of SARS-CoV-2 or SARS-CoV were loaded on Octet SA sensor and tested for real-time association and dissociation of the H014 Fab fragments. Global fit curves are shown as black dotted lines. The vertical dashed lines indicate the transition between association and disassociation phases.

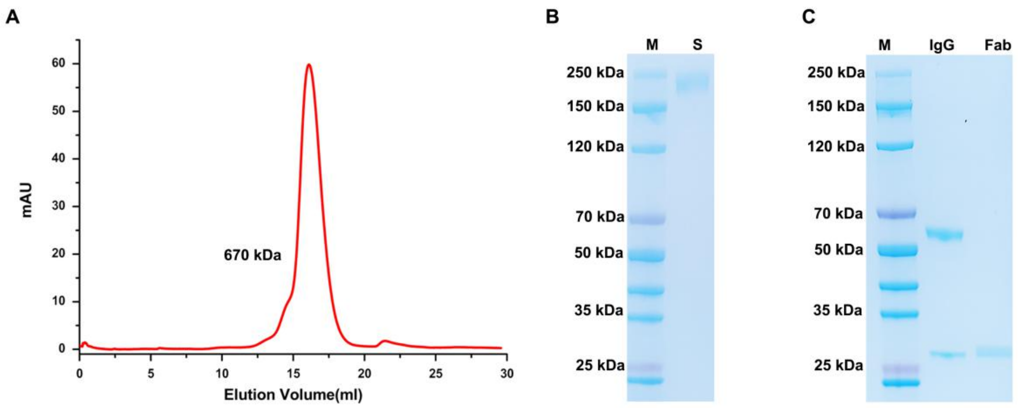


**Fig S3. Purification and characterization of SARS-CoV-2 S trimer and H014 Fab.** (**A**) Gel filtration profile of the affinity-purified SARS-CoV-2 S trimer. (**B**) SDS-PAGE analysis of the SARS-CoV-2 S trimer. (**C**) SDS-PAGE analysis of the H014 Ig G and its Fab fragment.

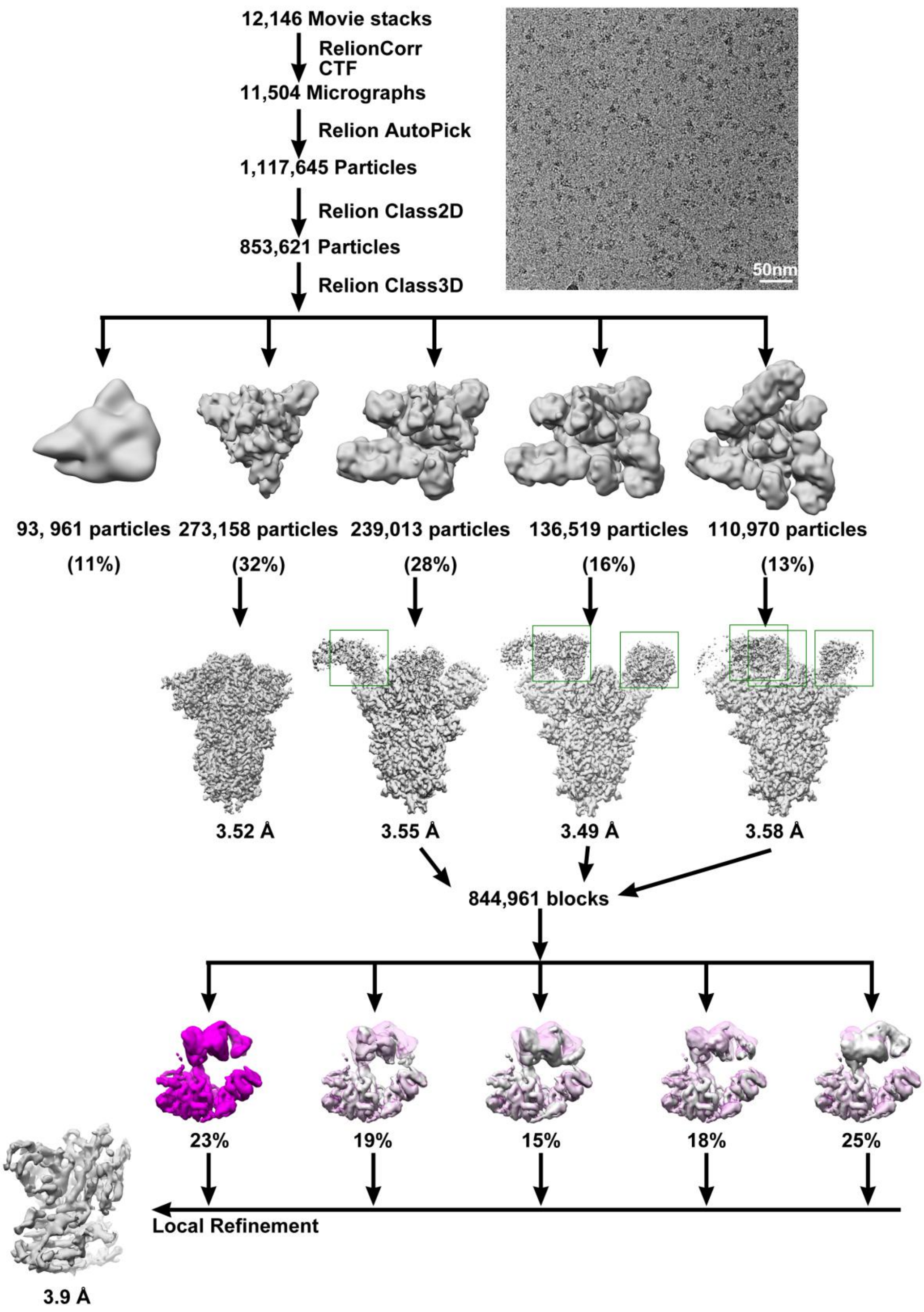


**Fig S4. Flow chart for Cryo-EM data processing.**

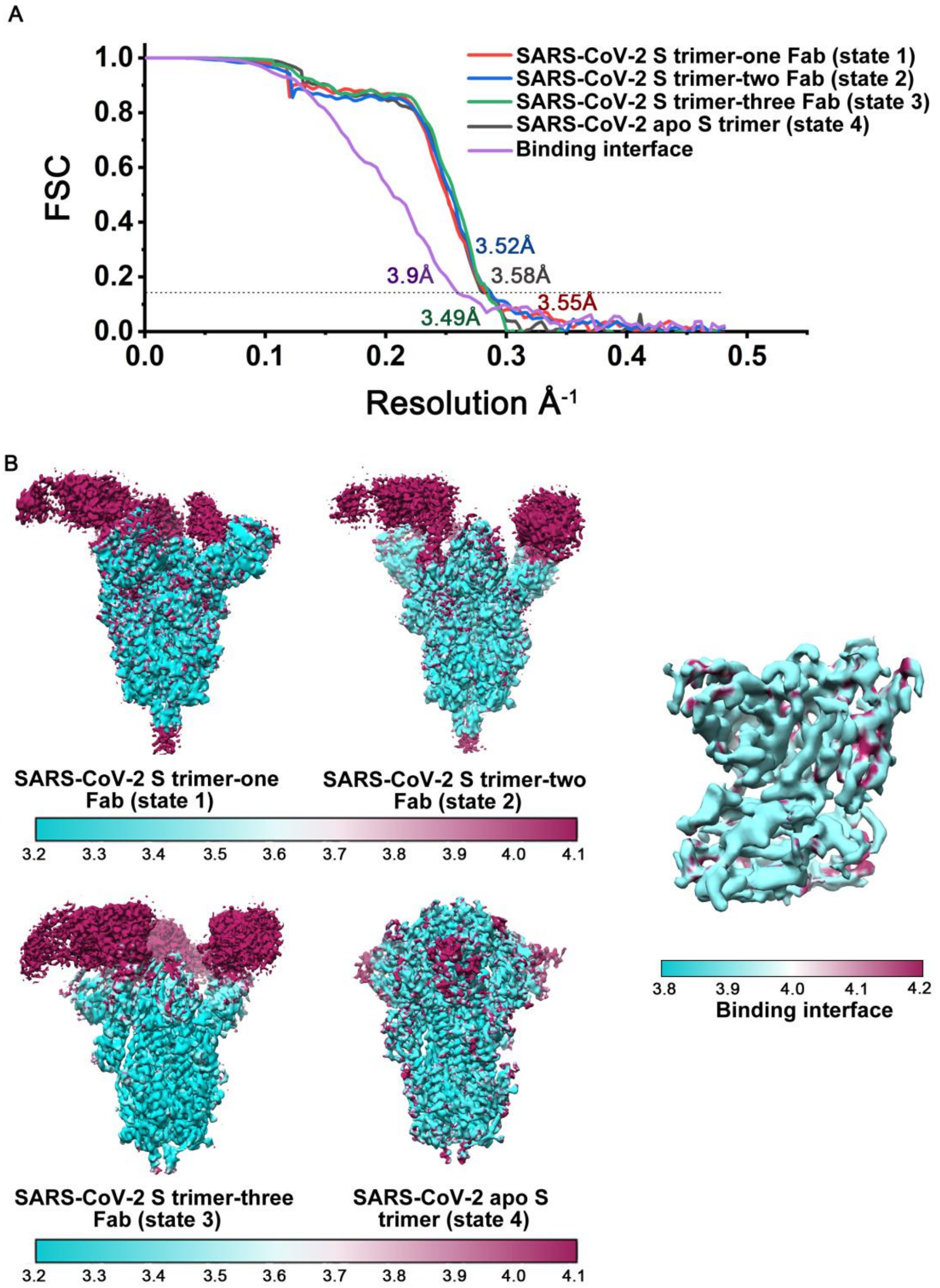


**Fig S5. Cryo-EM images and resolution evaluation of the EM maps of SARS-CoV-2 S trimer-H014 complexes.**

(**A**) The gold-standard FSC curves of the final maps. (**B**) Local resolution assessments of cryo-EM maps. Local-resolution evaluation of the maps of the *apo* SARS-CoV-2 S trimer, SARS-CoV-2 S trimer in complex with one Fab, SARS-CoV-2 S trimer in complex with two Fabs, SARS-CoV-2 S trimer in complex with three Fabs and binding interface using ResMap (*35*) are shown.

A

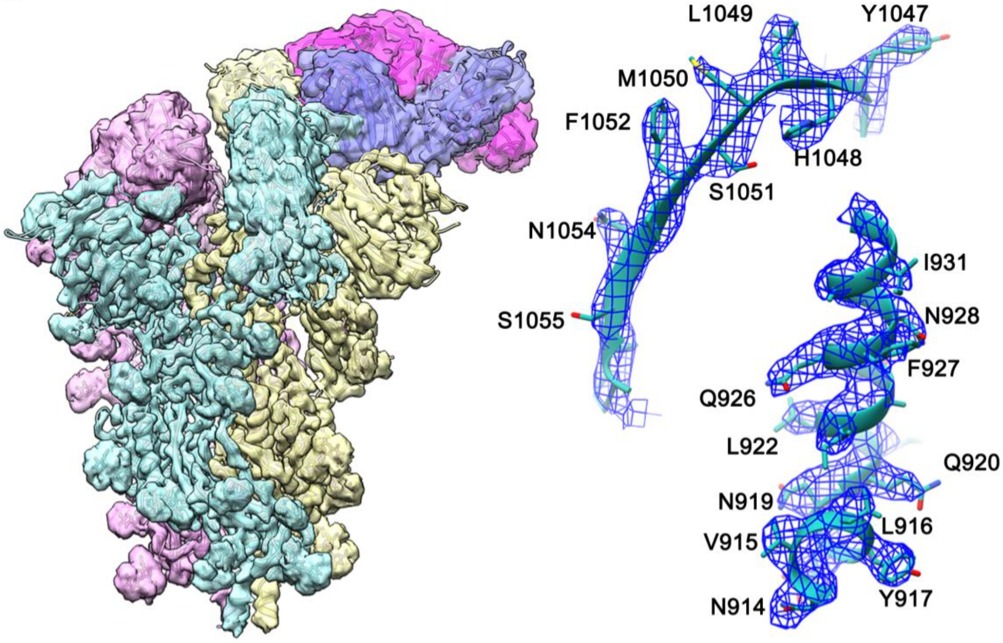


SARS-CoV-2 S trimer-one Fab (state 1)

B

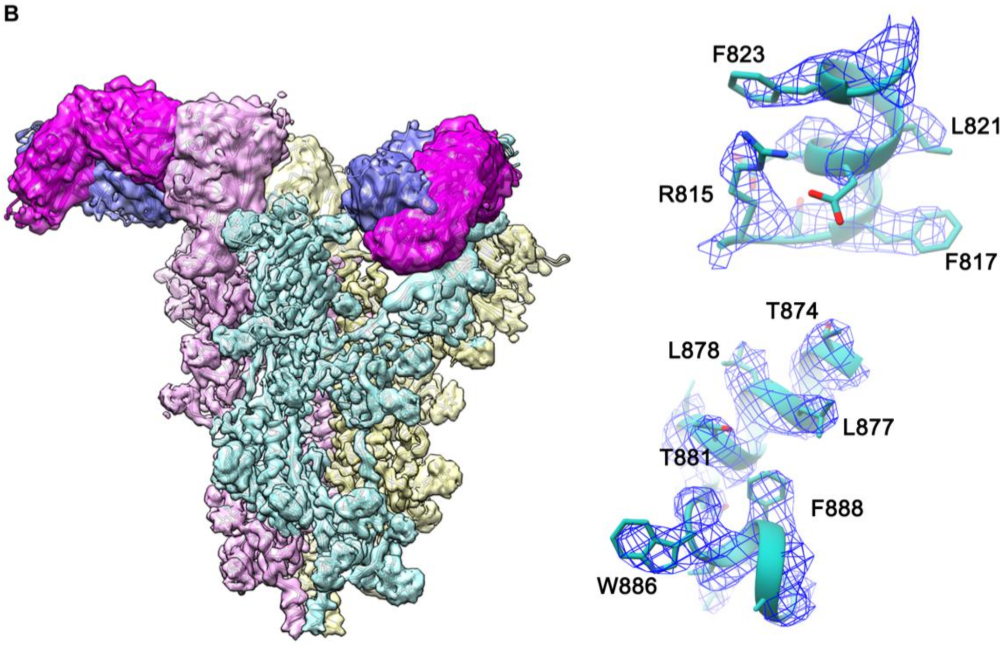


SARS-CoV-2 S trimer-two Fab (state 2)

C

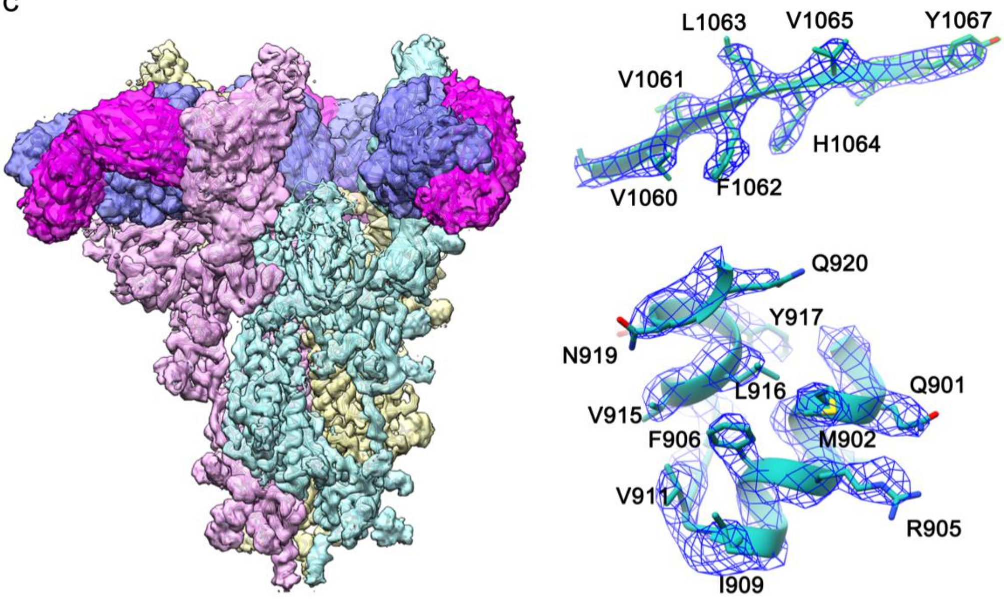


**SARS-CoV-2 S trimer-three Fab (state 3)**

D

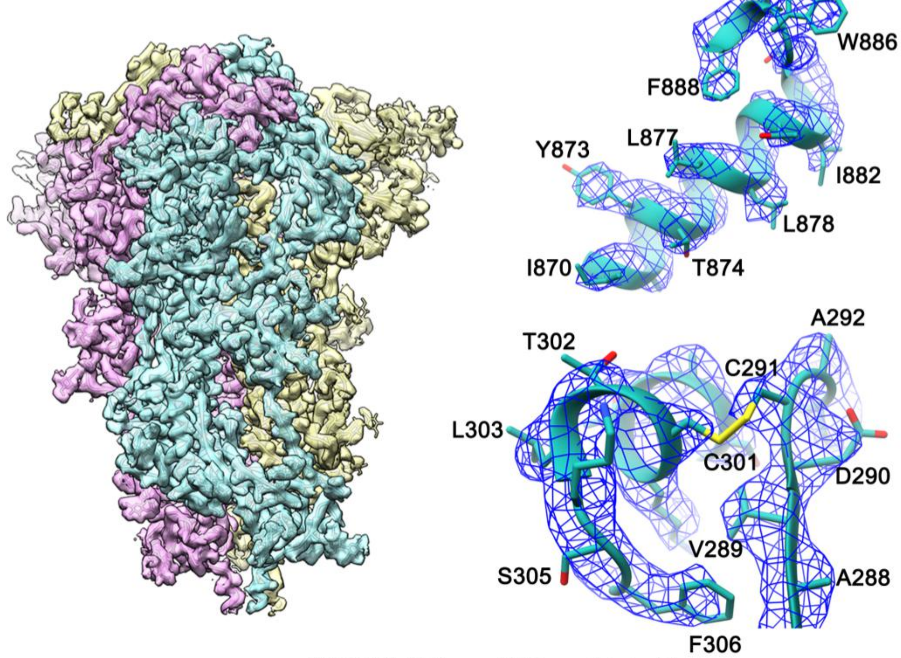


**SARS-CoV-2 apo S trimer (state 4)**

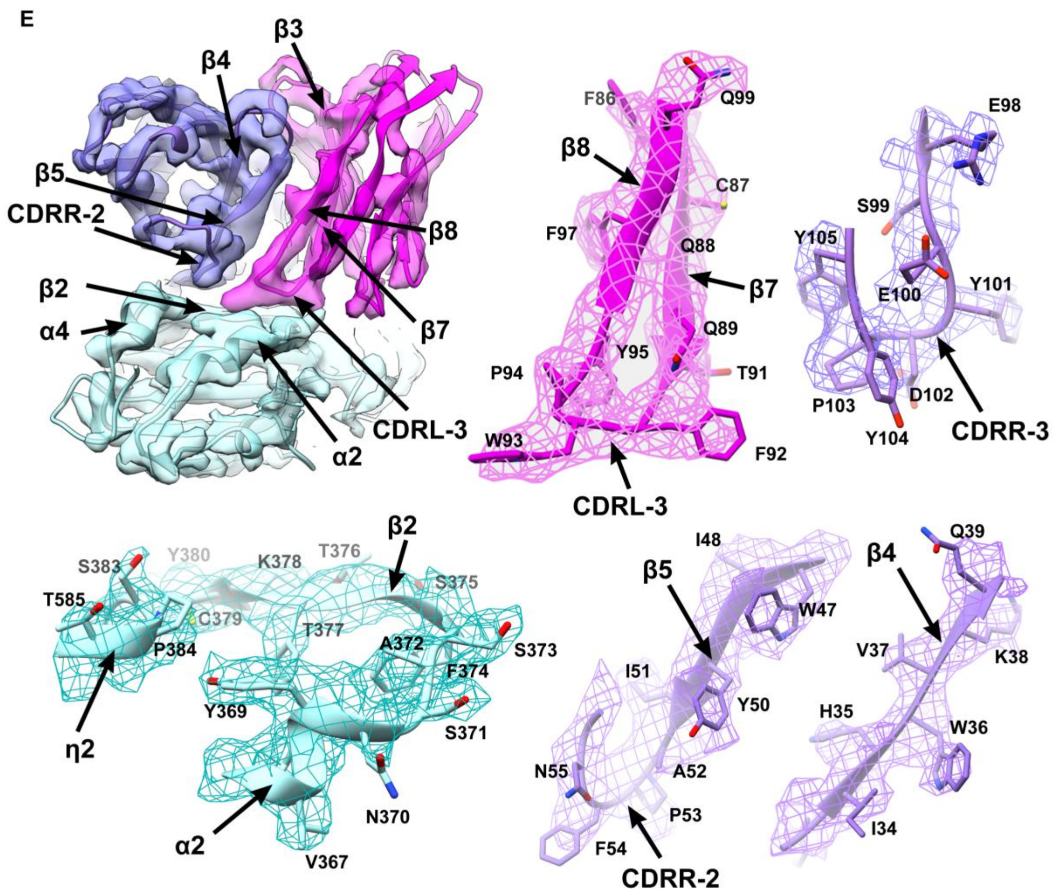


**Fig S6. Density maps and atomic models.**

Cryo-EM maps of SARS-CoV-2 S trimer in complex with one Fab **(A)**, SARS-CoV-2 S trimer in complex with two Fabs **(B)**, SARS-CoV-2 S trimer in complex with three Fabs **(C)**, the *apo* SARS-CoV-2 S trimer **(D)** and the binding interface **(E)** are shown. Color scheme is the same as in Fig. 2A. The enlarged panels show the density maps (mesh) and related atomic models. Residues are shown as sticks, oxygen atoms are colored red, nitrogens are colored blue and sulfurs shown in yellow.

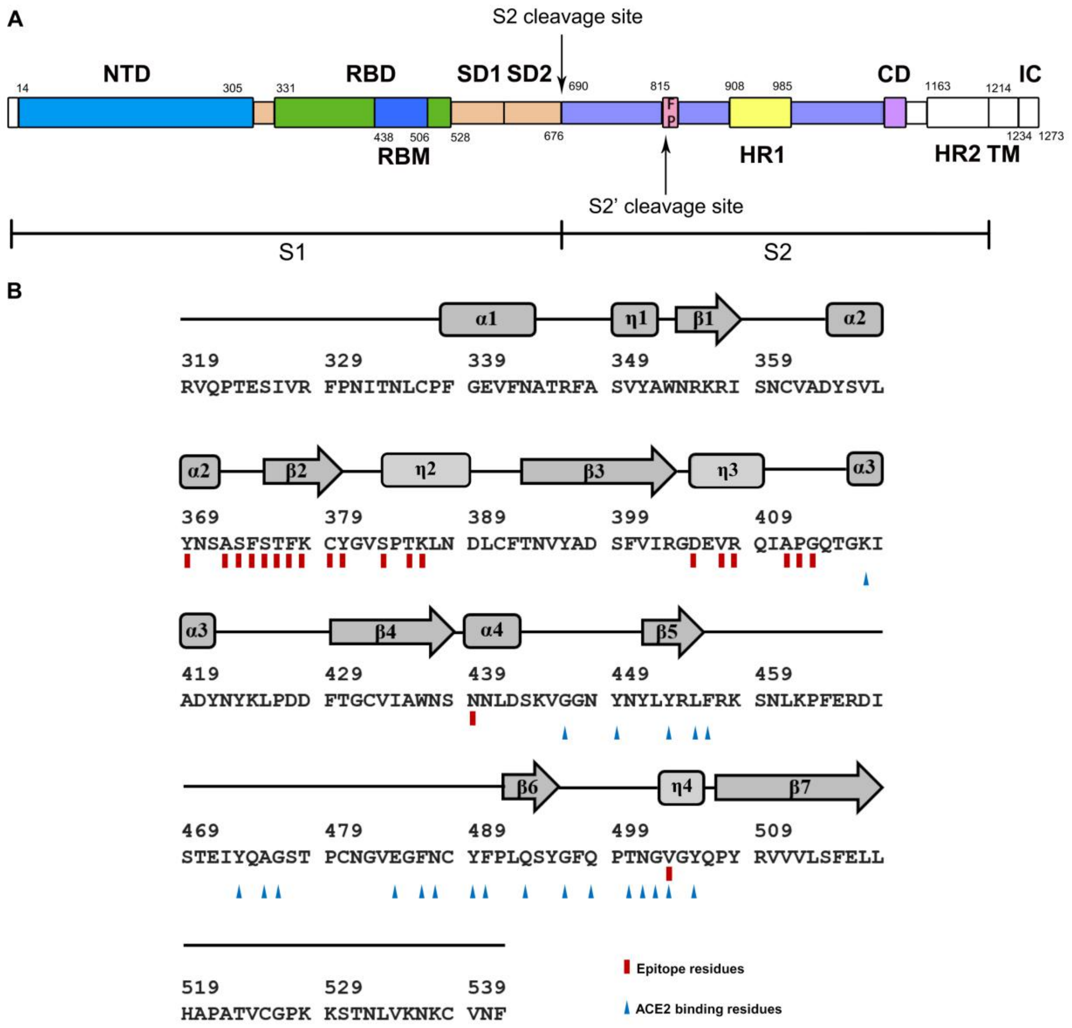


**Fig S7. Schematic diagram of SARS-CoV-2 S and secondary structure of the RBD.**

(**A**) Schematic showing the domain arrangement of the SARS-CoV-2 S monomer. NTD: N-terminal domain; RBD: receptor-binding domain; RBM: receptor-binding motif; SD1: subdomain 1; SD2: subdomain 2; FP: fusion peptide; HR1: heptad repeat 1; HR2: heptad repeat 2; TM: transmembrane region; IC: intracellular domain. (**B**) Primary sequence and secondary structural elements of SARS-CoV-2 RBD. The red rectangles and blue triangles indicate the residues in the SARS-CoV-2-S RBD that interact with H014 and ACE2, respectively.

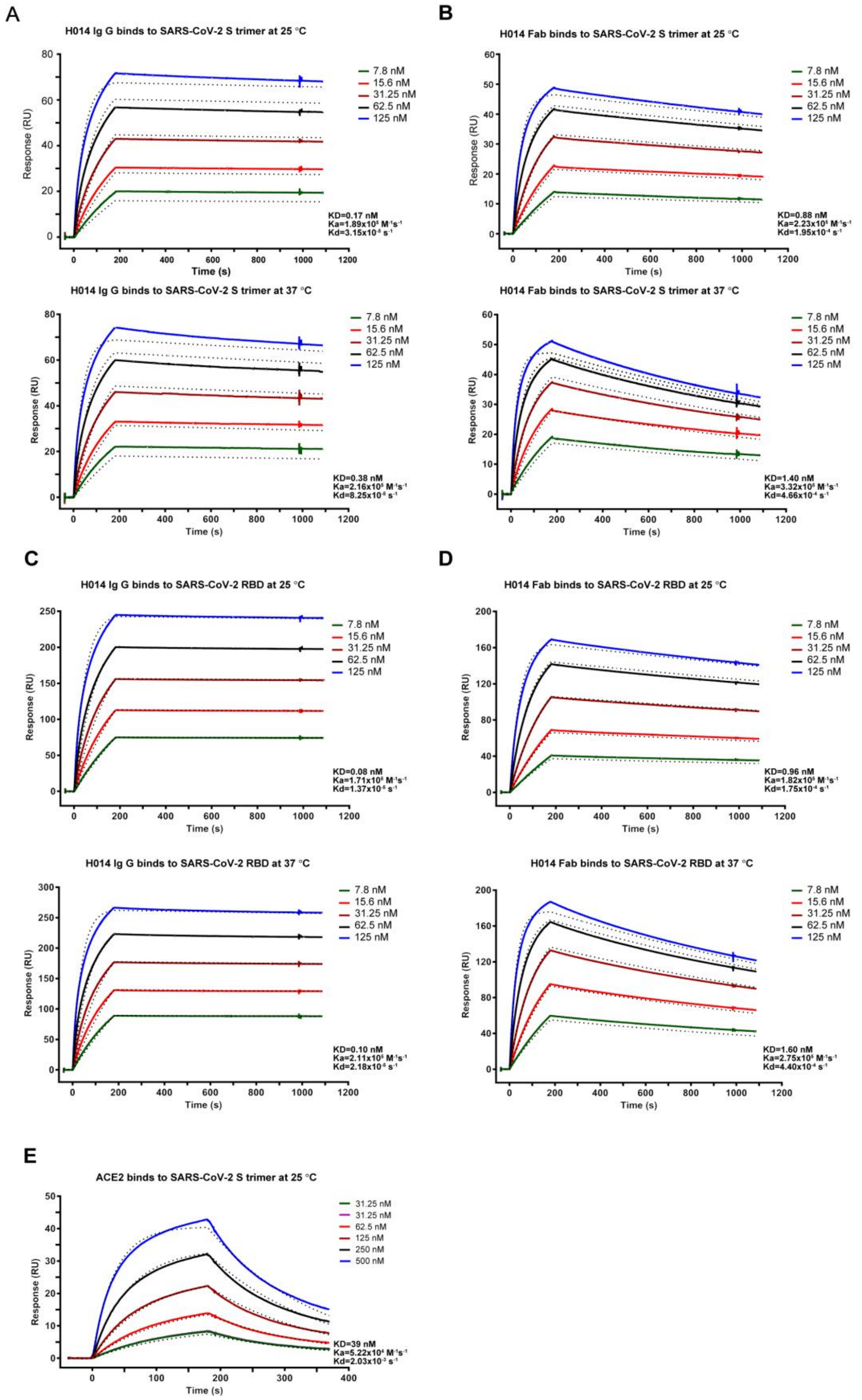


**Fig S8. BIAcore SPR kinetic profiles of H014/ACE2 and SARS-CoV-2 S trimer.** SARS-CoV-2 S trimer (**A** and **B**) was loaded onto the sensor. In the upper panel, binding of H014 Ig G (**A**) or H014 Fab (**B**) at various concentrations were monitored at 25 °C; in the bottom panel, binding of H014 Ig G (**A**) or H014 Fab (**B**) at various concentrations were monitored at 37 °C. SARS-CoV-2 RBD (**C** and **D**) was loaded onto the sensor. In the upper panel, binding of H014 Ig G (**C**) or H014 Fab (**D**) at various concentrations were monitored at 25 °C; in the bottom panel, binding of H014 Ig G (**C**) or H014 Fab (**D**) at various concentrations were monitored at 37 °C. (**E**) SARS-CoV-2 S trimer was loaded onto the sensor and binding of the ACE2 ectodomain at multiple concentrations were monitored at 25 °C. Global fit curves are shown as black dotted lines. The vertical dashed lines indicate the transition between association and disassociation phases.

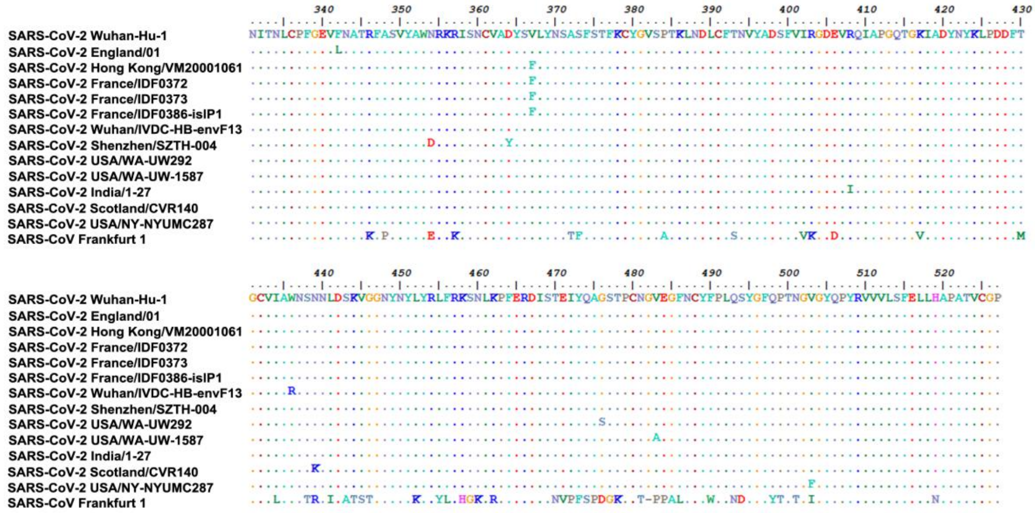


**Fig S9. Multiple sequence alignments of the mutations found in the RBDs of the currently circulating SARS-CoV-2 strains and SARS-CoV.**

The genome sequences used in alignments were downloaded from NCBI and GISAID with accession numbers: NC_045512.2, EPI_ISL_407071, EPI_ISL_412028, EPI_ISL_406596, EPI_ISL_406597, EPI_ISL_411219, EPI_ISL_408511, EPI_ISL_406595, EPI_ISL_418077, EPI_ISL_423014, EPI_ISL_413522, AY291315.1, EPI_ISL_425684 and EPI_ISL_428801 respectively. The alignments were analyzed by Clustal W (*41*) and BioEdit.

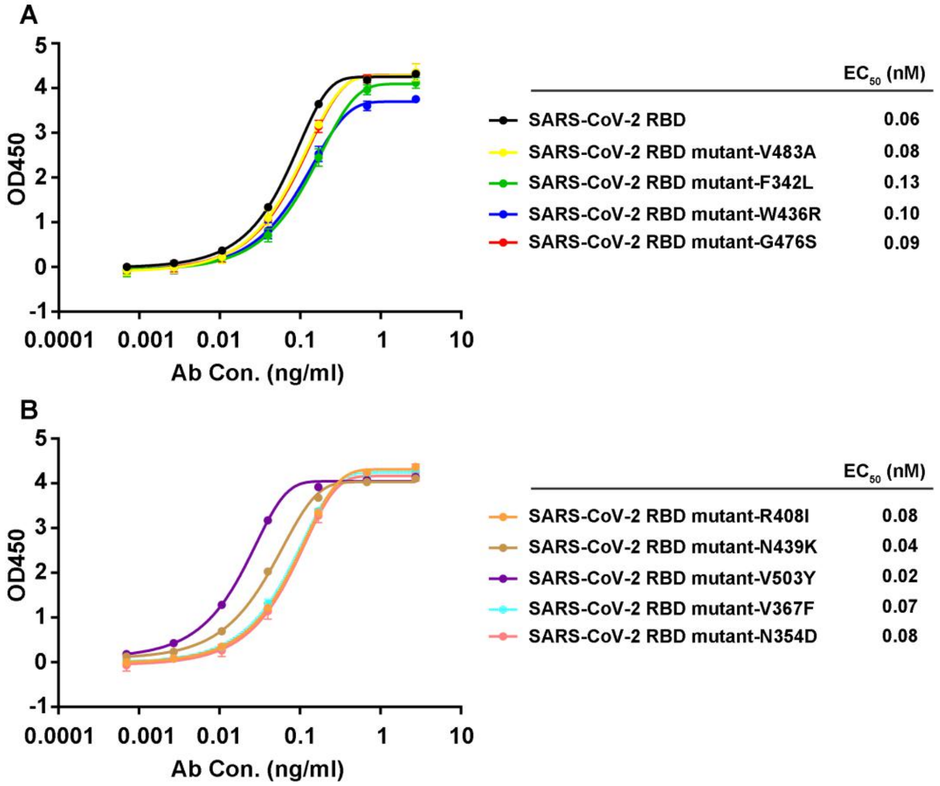


**Fig S10. Binding assays of SARS-CoV-2 RBD mutants to H014 by ELISA.**

Various SARS-CoV-2 RBD proteins harboring previously reported point mutations were analyzed for their binding abilities to H014 by ELISA.

### Table S1 Cryo-EM data collection and atomic models' refinement statistics

#### Data collection and reconstruction statistics

| Protein | State 1 | State 2 | State 3 | State 4 | Binding interface |
|---|---|---|---|---|---|
| Voltage (kV) | 300 | 300 | 300 | 300 | 300 |
| Detector | K2 | K2 | K2 | K2 | K2 |
| Pixel size (Å) | 1.04 | 1.04 | 1.04 | 1.04 | 1.04 |
| Electron dose ($e^-$/Å$^2$) | 60 | 60 | 60 | 60 | 60 |
| Defocus range (μm) | 1.25-2.7 | 1.25-2.7 | 1.25-2.7 | 1.25-2.7 | 1.25-2.7 |
| Final particles | 239，013 | 136，519 | 110,970 | 273,158 | 844,961 |
| Final resolution (Å) | 3.55 | 3.49 | 3.58 | 3.52 | 3.90 |

#### Models refinement and validation statistics

| | | | | | |
|---|---|---|---|---|---|
| Ramachandran statistics | | | | | |
| Favored (%) | 91.16 | 90.66 | 90.37 | 96.21 | 94.53 |
| Allowed (%) | 6.41 | 9.23 | 9.52 | 3.48 | 4.48 |
| Outliers (%) | 2.43 | 0.11 | 0.11 | 0.31 | 1.00 |
| Rotamer outliers (%) | 0.19 | 0.21 | 0.35 | 0.00 | 0.11 |
| R.m.s.d | | | | | |
| Bond lengths (Å) | 0.023 | 0.021 | 0.014 | 0.016 | 0.019 |
| Bond angles (°) | 1.043 | 1.511 | 1.016 | 1.051 | 1.332 |

**Table S2. Residues of H014 Fab fragment interacting with the SARS-CoV-2 S trimer at the binding interface (d < 4 Å)**

| S-RBD | | H014 Fab | |
|---|---|---|---|
| **Location** | **Residues** | **Light chain** | **Heavy chain** |
| **α2** | Y369 | W93 | |
| **α2-β2** | A372 | F92 | |
| | S373 | F92 | |
| | F374 | N91, F92, W93 | |
| | S375 | N91, F92, W93, Y95 | Y105 |
| **β2** | T376 | | Y105 |
| | F377 | W93 | S59 |
| | K378 | | Y50, S59, D102 |
| | C379 | | G56, T58 |
| | Y380 | | N55, G57 |
| **η2** | S383 | | K65 |
| | T385 | | L62, K65 |
| | K386 | | A67 |
| **η3** | D405 | | Y104 |
| | V407 | | D102 |
| | R408 | | N31, Y33, Y101, D102 |
| **η3-α3** | A411 | | N55 |
| | P412 | | N55 |
| | G413 | | F54 |
| **α3** | N439 | S29 | |
| **η4** | V503 | Y49 | |
